# How to train a post-processor for tandem mass spectrometry proteomics database search while maintaining control of the false discovery rate

**DOI:** 10.1101/2023.10.26.564068

**Authors:** Jack Freestone, Lukas Käll, William Stafford Noble, Uri Keich

**Affiliations:** School of Mathematics and Statistics F07, University of Sydney; Science for Life Laboratory, KTH Royal Institute of Technology, Stockholm, Sweden; Department of Genome Sciences, University of Washington; Paul G. Allen School of Computer Science and Engineering, University of Washington

**Keywords:** proteomics, false discovery rate control, tandem mass spectrometry

## Abstract

Decoy-based methods are a popular choice for the statistical validation of peptide detections in tandem mass spectrometry proteomics data. Such methods can achieve a substantial boost in statistical power when coupled with post-processors such as Percolator that use auxiliary features to learn a better-discriminating scoring function. However, we recently showed that Percolator can struggle to control the false discovery rate (FDR) when reporting the list of discovered peptides. To address this problem, we introduce Percolator-RESET, which is an adaptation of our recently developed RESET meta-procedure to the peptide detection problem. Specifically, Percolator-RESET fuses Percolator’s iterative SVM training procedure with RESET’s general framework of determining the list of reported discoveries in a target-decoy competition setup, where each putative discovery is augmented with a list of relevant features. Percolator-RESET operates in both a standard single-decoy mode and a two-decoy mode, the latter requiring the generation of two decoys per target. We demonstrate that Percolator-RESET controls the FDR in both modes, both theoretically and empirically, while typically reporting only a marginally smaller number of discoveries than Percolator in single-decoy mode. The two-decoy mode is marginally more powerful than both Percolator and the single-decoy mode and exhibits less variability than the latter.

## 1 Introduction

Tandem mass spectrometry provides a high-throughput method for detecting proteins in a complex sample. The approach involves digesting the proteins to peptides and then running a liquid chromatography tandem mass spectrometry (MS/MS) experiment to generate MS2 fragmentation spectra. Each of those spectra is then searched against a relevant (“target”) peptide database to find its best matching peptide, which combined with the spectrum defines the (optimal) peptide-spectrum match (PSM). The PSMs can then be used as evidence to deduce the presence of relevant peptides and proteins in the sample.

In practice a PSM can be incorrect, meaning that the matched peptide did not generate the corresponding spectrum. It follows that a false discovery can be made at any of the three levels of the analysis: spectrum identification, as well as peptide and protein detection. The canonical approach to the statistical validation of the reported PSM, peptide, or protein discoveries is through controlling the false discovery rate (FDR) via target-decoy competition (TDC) [4].

TDC originated in the mass spectrometry community; however, the underlying competition-based approach to controlling the FDR has more recently been popularized in the context of feature selection in the statistics and machine learning communities [1]. The overarching theme connecting these applications is that of hypothesis testing, where we define a set of null hypotheses and the goal is to reject a subset of these hypotheses while controlling the FDR. In the MS/MS setup a null hypothesis can be “the considered PSM is incorrect,” or alternatively “the considered peptide or protein is not present in the sample,” and in variable selection in regression it is “the considered variable is not included in the model” (e.g., its coefficient is zero in a linear model).

A valid application of TDC hinges on our ability to generate for each null hypothesis two scores: a target score that summarizes our evidence against the null, and a decoy score that is independently drawn assuming that the null is correct. For example, if the null is that the PSM is incorrect, then the target score is the search tool’s original target PSM score, and the decoy score is obtained from the corresponding decoy PSM generated by searching the same spectrum against a database of decoy peptides. In this setting, decoys are typically constructed by reversing or randomly shuffling the original target peptides.

Notably, if the null hypothesis is true, then both the decoy and the target scores are sampled from the same null distribution. Therefore, each one is equally likely to win the head-to-head competition between the two scores to define the corresponding winning score. This allows us to associate with each null hypothesis the winning score as well as a target/decoy label indicating which of the two was the winner. Therefore, if we sort the winning scores in decreasing order, then the number of decoy wins at any score threshold can be used to estimate the number of incorrect target wins, or the number of false discoveries. Hence, we can estimate the FDR by dividing this number of decoy wins by the corresponding number of target wins above the same score threshold. Moreover, to control the FDR at, say 1%, we can look for the smallest score threshold such that the estimated FDR (slightly modified by adding one “pseudo” decoy win to the numerator) is still ≤ 1%. Notably, it has been established that, assuming that a true null’s target and decoy scores are independent of all other scores then this TDC procedure theoretically guarantees FDR control [1, 10].

Unfortunately, applying TDC to database search results can become more complicated when those results are first post-processed by a machine learning algorithm that takes advantage of features such as the peptide’s length or its presumed charge state to re-rank the PSMs. The general idea was pioneered by the PeptideProphet algorithm [12], which involved training a linear discriminant analysis (LDA) model to classify PSMs on the basis of a hand-curated set of “correct” and “incorrect” PSMs. This approach suffered from poor generalizability when the data used to train the model had different characteristics than the data being classified. This problem led to the development of Percolator [11], which is a semi-supervised machine learning algorithm that trains its model using a collection of target and decoy PSMs. The idea is that the decoy PSMs can be labeled “incorrect,” whereas the targets are a mixture of “correct” and “incorrect” PSMs. Importantly, because target and decoy PSMs can be generated automatically, no hand curation of labels is necessary, and Percolator can be retrained for every new dataset. This approach guarantees that the training set and test set distributions are similar. The semi-supervised approach has become the *de facto* standard in the field.

Despite its popularity, we have recently shown that in practice Percolator can fail to control the FDR [7]. The problem arises because Percolator uses decoys in two fundamentally different ways: first to train a machine learning model to distinguish between correct and incorrect PSMs, and second to estimate FDR using the TDC procedure. Recognizing this challenge, Percolator uses a cross-validation procedure to attempt to prevent information about the target/decoy labels from leaking from the training set to the test set [9]. However, we argued that in practice Percolator’s training process can indirectly peek at the target/decoy labels of the test set, and hence the overall procedure may fail to control the FDR.

Specifically, we highlighted the problem of “multiplicity” where the same analyte creates multiple spectra — a phenomenon that is essentially guaranteed when multiple runs are combined. These multiple spectra can create nearly identical PSMs that are shared among the training and testing folds, thereby undermining Percolator’s cross-validation mechanism [7]. While this multiplicity problem can be solved if Percolator switches from PSM-level to peptide-level training, the question of whether Percolator’s cross-validation scheme rigorously controls the FDR remains open. This motivated our recent introduction of “RESET” (REScoring via Estimating and Training) [5]. RESET is a meta-procedure designed to take advantage of side-information, or additional features, such as those used by Percolator, so as to improve the ranking of the correct discoveries while controlling the FDR in the competition setup. RESET achieves this goal by first randomly splitting the winning decoys into two sets of roughly similar size: a training set and an estimating set. It then trains a classifier that has access to the primary score and all the additional features to distinguish between the training decoys and the combined set of estimating decoys and target wins. Notably, a variety of classifiers can be used at this step but, critically, the classifier should not have access to the label that distinguishes between the target wins and the estimating decoys, which we jointly refer to as “pseudo-targets.” Therefore, after reranking the pseudo-targets by the newly learned score, the number of (estimating) decoy wins at any given score threshold can still be used to estimate the number of false target discoveries (although in this case we need to multiply the number of estimating decoys by two).

Here we introduce “Percolator-RESET”, which is an of adaptation of RESET to the peptide detection problem — the goal of identifying which peptides are present in the sample. Percolator-RESET borrows from Percolator its iterative training of an SVM that aims to distinguish between correct and incorrect discoveries. We argue that under assumptions that naturally extend those that TDC relies on, Percolator-RESET rigorously control the FDR, and we augment this theoretical analysis by empirically demonstrating its robust control of the FDR in a setup where Percolator’s control is questionable. Comparing the power (i.e., the number of discoveries) of two tools we find that, in spite of its much tighter FDR control, Percolator-RESET gives up very little to Percolator. Moreover, Percolator-RESET can take advantage of a second randomly drawn decoy database, which delivers slightly more discoveries than Percolator (as well as more than its single-decoy version). At the same time, using the additional decoy database reduces Percolator-RESET’s variability, which is inherent to its random split of the decoy set.

In addition to giving the user the option of using a second decoy database, Percolator-RESET also provides the option of using peptide-level target-decoy pairing information. This is motivated by our earlier analysis showing that using this pairing to conduct a second, peptide-level competition delivers more discovered peptides than when using only the PSM-level competition [16]. While potentially beneficial, many search tools do not provide the user with this target-decoy pairing information, and moreover they often integrate a decoy construction procedure that prevents any natural pairing. For example, the most common decoy construction is the reversal of the entire target protein before it is digested into decoy peptides. These decoy peptides are different from ones created by reversing the target peptides directly and, in particular, their mass often differs slightly from that of the corresponding reversed peptide making for a somewhat unnatural pairing. This problem becomes much more pronounced when non-tryptic digestion is considered. To make Percolator-RESET compatible with search methods that do not provide pairing information, the method was designed to work in the absence of pairing information while still rigorously controlling the FDR.

An open source, Apache licensed Python implementation of Percolator-RESET is available at https://github.com/freejstone/Percolator-RESET. The unpaired single-decoy option will also be implemented as part of an upcoming release of Percolator (v4.0).

## 2 Methods

### 2.1 Percolator-RESET overview

Percolator-RESET is an adaptation of our RESET meta-procedure (Algorithm 1) to the problem of peptide detection: given MS/MS data, report a list of peptides that are present in the sample while controlling the FDR. We begin by outlining Percolator-RESET’s main steps, which we subsequently expand on.

1. Use the target and decoy PSMs generated by the search tool to define the list of considered target and decoy peptides and their scores while taking into account whether
  - one or two decoy databases were searched,
  - the decoy peptides are paired (in the single-decoy case) or matched (in the two-decoy case) with the target peptides, and
  - variable modifications were considered.
2. Apply Percolator-RESET’s core of rescoring the considered peptides using the following strategy:
  - Randomly assign each decoy peptide as an *estimating decoy* or a *training decoy*.
  - Define the *pseudo-targets* as the combined set of target and estimating decoy peptides.
  - Use the peptide scores and the additional features to train an initial SVM to distinguish between a positive and a negative set of peptides where:
    - the (fixed) negative set is the set of training decoys, and
    - the positive set is the set of high-scoring pseudo-targets selected by controlling the FDR using TDC applied to the training decoys (as decoys) and the pseudo-targets (as targets).
  - Iteratively use the trained SVM to rescore the considered peptides and repeat the last step to train a new SVM using the new scores. Note that the positive set typically changes with each iteration.
  - After a fixed number of iterations, return the last-trained-SVM scores of all considered peptides, as well as the selected set of estimating decoys.
3. Use TDC to report the discovered target peptides while controlling the FDR using the estimating decoy peptides. This steps takes into account whether a single or two decoy databases were searched.
4. Optionally, report a list of high-scoring PSMs associated with the reported peptides.

### 2.2 Target-decoy competition

Unless otherwise stated, when we mention “TDC” we specifically refer to its decoy-estimated FDR controlling component, which we formalize here as `Selective SeqStep` / `Selective SeqStep+`, or `SSS` / `SSS+` for short (Algorithm 2). Note that these procedures are essentially the same-named procedures described by Barber and Candès [1], and they generalize the canonical TDC by allowing arbitrary ratios of targets to decoys.

Specifically, canonically in TDC a true null hypothesis, say an incorrect PSM, is equally likely to be a target or a decoy, which corresponds to setting *c* = 1*/*2 in Algorithm 2. Indeed, in this case we multiply the number of decoys by *c/*(1 − *c*) = 1, reflecting the fact that, on average, for every decoy there is one incorrect target. By varying *c*, these procedures allow us to account for the case where the decoy peptide database is, say, half the size of the target’s. Indeed, in this case we would set *c* = 2*/*3, or *c/*(1 − *c*) = 2, thus taking into account that, on average, for every decoy there are two incorrect targets.

The difference between the `SSS` and `SSS+` is whether the +1 pseudo-count is added to the number of decoys when estimating the FDR. This +1 is required to guarantee finite sample control of the FDR [1].

### 2.3 Percolator-RESET

#### 2.3.1 Defining the considered peptides and their scores

Percolator-RESET starts by executing a PSM-level competition, where for each spectrum *σ* only its best matching peptide across the target and the one or two decoy databases is retained. “Best” here is in terms of the search engine’s primary score, where in our experiments we used the Tailor similarity score [19] when processing Tide’s searches [3], and the Hyperscore when processing MSFragger’s [15].

In practice, this competition was implemented somewhat differently with Tide and MSFragger, especially when using two decoys. Specifically, in MSFragger’s case we simply searched a concatenated target-decoy `.fasta` file, where each target protein in the file was accompanied by two decoy proteins that were constructed as described in Section 2.5 below. In Tide’s case we applied `tide-search` twice, once for each of the two randomly generated peptide-shuffled decoy databases produced by `tide-index`. The best target match for a given spectrum does not vary between the two searches, but that optimal target PSM will typically have different Tailor scores across the two searches. This is because the Tailor score normalizes the raw PSM score with respect to the quantiles of all the target and decoy matches to the given spectrum, and the decoys vary between the two searches [19]. We addressed this variation by averaging the two Tailor scores assigned for each target PSM before competing it with the two decoy PSMs associated with the same spectrum. We next define each peptide’s score as the maximal PSM score associated with that peptide, and we let the peptide inherit all the features associated with this maximal scoring PSM (charge, peptide length, delta mass, etc.). In the absence of target-decoy peptide pairing information (note that “pairing” should be construed as “matching” in the two-decoy mode), the list of considered peptides consists of all those which are associated with at least one PSM. If, however, we have such pairing information then following the PSM-and-peptide protocol of [16], for each paired target-decoy peptides we only add the higher scoring one to the list of considered peptides. Similarly, in the two-decoy application we use the potential availability of the analogous matching information to, again, retain only the top scoring peptide from every triplet of a target and its two matched decoys. Algorithm 3a summarizes the procedure described so far for the two-decoy application, where the adjustments for the single-decoy version are straightforward.

We recently pointed out that if variable modifications are specified, then the decoy database can no longer provide adequate competition against some of the incorrectly identified target peptides [6]. Specifically, TDC relies on the assumption that a false discovery is equally likely to be a target or a decoy. However, when variable modifications are specified the target database will often contain clusters of highly similar peptides corresponding to variable modifications of the same unmodified (“stem”) form of a single peptide. The problem is that the canonically constructed decoys (obtained by shuffling or reversing the target peptides) cannot account for mismatches within such clusters. To address this problem we employ the dynamic-level competition introduced in [6]: when variable modifications are considered, we switch from peptide-level to stem-level analysis, where we cluster the PSM-associated peptides according to their stem, or unmodified residue sequence. For example, we would cluster the peptides P[10]EPTIDE, PEP[10]TIDE, P[10]EP[10]TIDE, as well as any other peptidoform of PEPTIDE that has an associated PSM. We then select from each cluster its highest scoring peptide as the cluster’s representative.

In the absence of target-decoy pairing information, the considered peptides are the above selected representatives. However, given such pairing information we can readily extend it to the clusters, and hence to the clusters’ representatives. Therefore, given the extra information we execute our optional second target-decoy competition at the representative level, keeping only the higher-scoring of each competing pair (or triplet in the two-decoy mode) of representatives, together with its associated features. Note that in this case, the competing representatives are paired at the cluster level, and each of them might be paired with other peptides from the cluster of the competing representative. Algorithm 3b summarizes this procedure for selecting the representative peptides in the two-decoy with variable modification application. Again, the adjustments for the single-decoy version are straightforward.

#### 2.3.2 Using additional features to learn a better scoring scheme

Percolator-RESET’s approach to learning a better scoring function follows Percolator’s approach with some notable changes that we highlight below. Recall that training an SVM classifier requires a negative and a positive set. To define these, we first independently randomly assign each considered decoy peptide — as defined in the previous section — as a *training decoy* or an *estimating decoy* with equal probability of 1/2. We refer to the combined list of estimating decoys and considered target peptides as *pseudo-targets*, alluding to the fact that the SVM jointly consider them as targets. The fixed negative set consists of all the training decoys, while the positive set is iteratively updated by applying TDC to select a list of pseudo target discoveries while controlling the FDR at level *α*_1_. This training-related FDR threshold is independent of the overall FDR threshold *α*, and we use the same *α*_1_ = 0.01 as in Percolator.

Percolator-RESET introduces a couple of major changes to Percolator’s definition of the positive and negative sets. First, it applies TDC at the peptide level, whereas Percolator does it at the PSM level. Second, it uses roughly half the decoys as the negative set, and its positive set includes a subset of the pseudo-targets. In contrast, Percolator uses a three-fold cross-validation scheme, so it uses all the decoys in the two training folds, and Percolator’s positive set is a subset of the targets in the same two folds.

Because, when defining the positive set, Percolator-RESET considers the estimating decoys as part of the pseudo-targets, the TDC procedure is adjusted accordingly. Specifically, in the single-decoy case we use `SSS+` with *c* = 3*/*4, or *c/*(1 − *c*) = 3 because, on average, for each training decoy there are three incorrect pseudo-targets (one estimating decoy and two incorrect targets). In practice, this means that number of training decoys (plus 1) is multiplied by 3. In the two-decoy case, because of the inherent 2:1 ratio of decoys to targets, for each training decoy there are two incorrect pseudo-targets (one estimating decoy and one incorrect target), so in this case we use *c* = 2*/*3, or *c/*(1 − *c*) = 2.

If the dataset is sufficiently small, then it is possible that the positive training set as defined above will be empty. In this case, we first repeat the procedure but using `SSS` rather than `SSS+` in Algorithm 2 (i.e., we remove the +1 pseudo-count). If the newly defined positive set is still empty, then we iteratively increase the training FDR threshold from its initial value of *α*_1_ = 0.01 by increments of 0.005 until `SSS` returns a non-empty discovery set. This set is then used as the positive training set.

Using the positive and negative training sets, and their associated features (including the scores), we train an SVM to rescore each considered peptide, and this process is repeated a fixed number of iterations (we used `total iter` = 5). Note that initially, the scores used by TDC in defining the positive set are those assigned by the search engine, but subsequent iterations use the previously trained SVM to rescore the considered peptides before applying TDC. An algorithmic description of the procedure is given in Algorithm 4.

#### 2.3.3 SVM training in Percolator-RESET

Because the positive and negative training sets are expected to be unbalanced, we select the class weights *C*^+^, *C*^*−*^ ∈ {0.1, 1, 10} using the following three-fold cross validation procedure, which is similar but not identical to Percolator’s:

1. Randomly split the union of the positive and negative sets defined in the previous section into three folds.
2. Holding out one fold at a time, do:
  a. For each pair of considered weights, (*C*^+^, *C*^*−*^), train the SVM on the positive and negative subsets of the other two folds.
  b. Apply the learned SVM weights to the held-out fold.
  c. Apply `SSS+` to the rescored, held-out fold using the positive subset as targets and the negative subset as decoys with *α*_1_ = 0.01 (and *c* = 3*/*4 or *c* = 2*/*3 depending on whether one or two decoys are used).
  d. Add the resulting number of positive (pseudo-target) discoveries to the tally of the considered (*C*^+^, *C*^*−*^) pair.
3. Choose the (*C*^+^, *C*^*−*^) pair that has the largest number of discoveries. Ties are broken arbitrarily according to the Python function `GridSearch`.
4. If that maximal number of discoveries is 0, then repeat the cross-validation process, first using `SSS` instead of `SSS+`, then gradually increasing *α*_1_ exactly as described above when defining the positive set, until that maximal number of discoveries is finally positive.

#### 2.3.4 Selecting the reported peptides using TDC

After rescoring the considered peptides, the training decoy peptides are removed and TDC is applied to the remaining set of estimating decoys (now revealed as such) and targets. In the single-decoy application, for every estimating decoy there are on average two incorrect targets (because we used half of the decoys as training decoys). Therefore, to control the FDR among the list of reported target peptides in this case we apply `SSS+` with *c* = 2*/*3, or *c/*(1− *c*) = 2. Similarly in the two-decoy application, with the training decoys thrown out we expect a ratio of 1:1 between the estimating decoys and incorrect targets; hence, in this case we apply `SSS+` with the TDC-canonical *c* = 1*/*2, or *c/*(1 − *c*) = 1.

#### 2.3.5 Optionally reporting PSMs

Percolator-RESET promises theoretical FDR control only at the (representative) peptide level, because the PSM level suffers from a multiplicity issue that violates the usual independence assumptions [10, 16]. Understandably, some sort of PSM-level analysis is often desirable. For example, when using an open search, multiple matches to the same stem peptide may correspond to distinct types of modifications of that peptide. Hence, similar to CONGA [6], we additionally report a list of PSMs that are associated with the originally discovered peptides. The process is essentially the same as the one used in CONGA: we first define a set of mass bins for each discovered (stem) peptide by a greedy clustering algorithm that takes into account the specified isolation window (Supplementary Algorithm 5). Then the highest scoring of all the PSMs with the same peptide that fall into the same mass bin is reported. Highest here is judged by the original peptide score (e.g., Tailor or Hyperscore) instead of the SVM score, since the alternative may lead to a PSM outscoring the original PSM that was responsible for discovering the peptide in the first place. The list of PSMs is enhanced with a label for each PSM, which indicates whether the PSM scored above the SVM score threshold, giving the user additional confidence in those PSMs.

### 2.4 Entrapment setup

Our entrapment experiments followed the same setup as described in [6, Section 3: Entrapment run searches] with some minor variations listed below. In brief, we used a dataset generated from the standard protein mix database, ISB18 [14]. The combined target databases consisted of increasing fractions of the original ISB18 peptides — the in-sample proportion of the randomly selected subset of ISB18 peptides was increased from 25% to 100% — combined with a fixed entrapment database from the castor plant proteome. As noted in [6, Table 1], the entrapment-to-in-sample peptides ratio was at least as high as 636:1. This allows us to use the lower-bound method [20, Equation (2)] for providing evidence of both FDR-control or its lack thereof, because the difference between the combined [20, Equation (1)] and the lower-bound methods is negligible in this case.

**Table 1:**
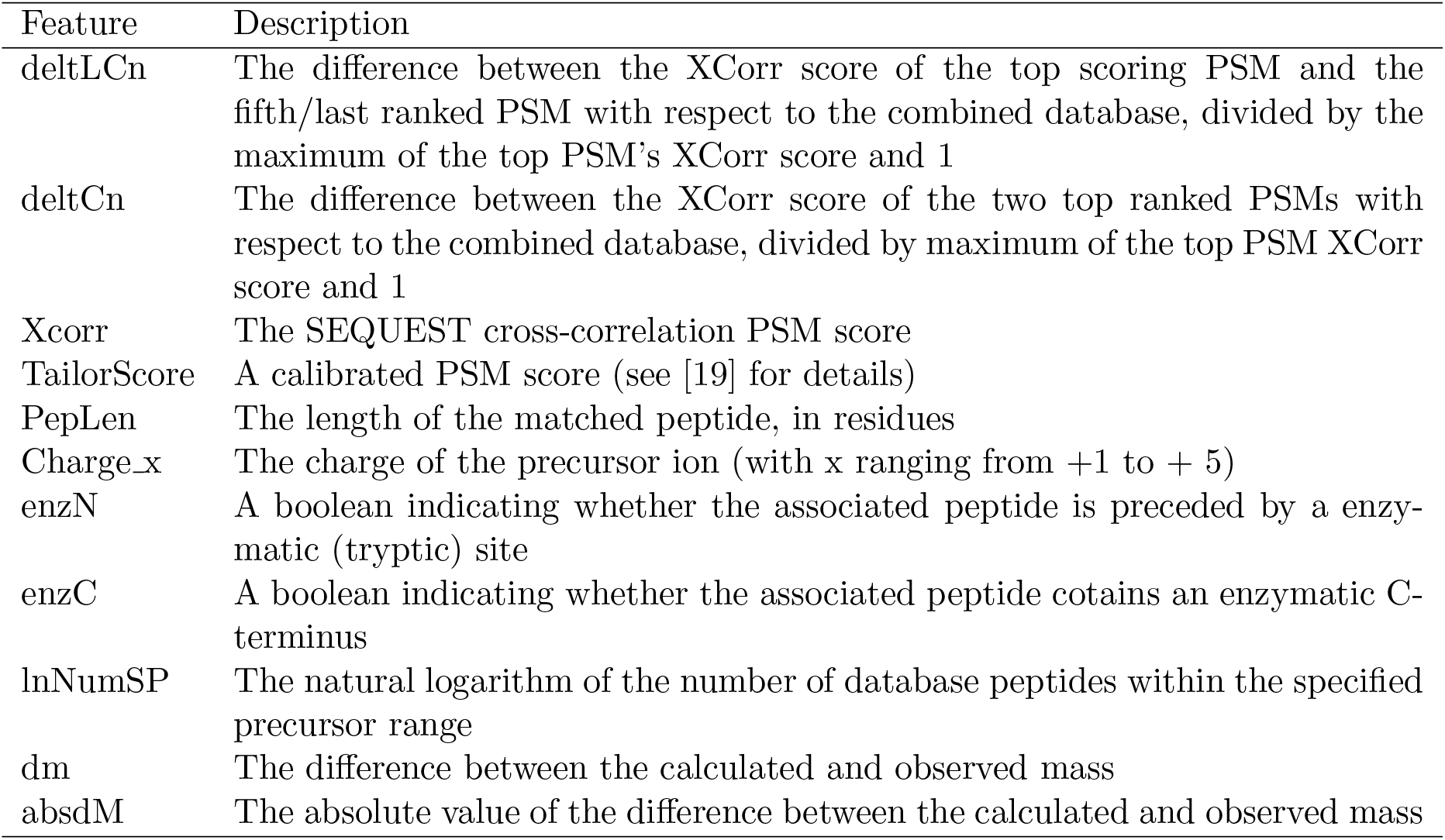
List of described features used in Tide. The list of features considered when applying Percolator-RESET, adapted from https://crux.ms/file-formats/features.html.

| Feature | Description |
| --- | --- |
| deltLCn | The difference between the XCorr score of the top scoring PSM and the fifth/last ranked PSM with respect to the combined database, divided by the maximum of the top PSM's XCorr score and 1 |
| deltCn | The difference between the XCorr score of the two top ranked PSMs with respect to the combined database, divided by maximum of the top PSM XCorr score and 1 |
| Xcorr | The SEQUEST cross-correlation PSM score |
| TailorScore | A calibrated PSM score (see [19] for details) |
| PepLen | The length of the matched peptide, in residues |
| Charge_x | The charge of the precursor ion (with x ranging from +1 to + 5) |
| enzN | A boolean indicating whether the associated peptide is preceded by a enzymatic (tryptic) site |
| enzC | A boolean indicating whether the associated peptide contains an enzymatic C-terminus |
| lnNumSP | The natural logarithm of the number of database peptides within the specified precursor range |
| dm | The difference between the calculated and observed mass |
| absdM | The absolute value of the difference between the calculated and observed mass |

The variations on the protocol described in Freestone *et al*. [6] were as follows:

- In empirically estimating the FDR at any given threshold we averaged the estimated false discovery proportions (FDPs) over 100 runs (instead of over 20 previously), each using a different randomly generated decoy database.
- The searches conducted by Tide in open and narrow search modes used an updated version of Crux (v4.1.6809338 [13]). The same version was also used for all other analyses using the Crux toolkit that are reported in this paper.
- The option `--concat` in `tide-search` was set to `T` for searches subsequently used by Percolator and by the single-decoy mode of Percolator-RESET, while it was set to `F` for searches used in the two-decoy mode of Percolator-RESET.
- Percolator-RESET was employed in both single-decoy and two-decoy modes, as well with pairing information (PSM-and-peptide) and without (PSM-only).

### 2.5 Searches of the PRIDE-20 dataset

We used the same PRIDE-20 spectrum files and database FASTA files as described in [6, Section 3: PRIDE-20 searches]. Briefly, this is a collection of publicly available MS/MS datasets from 20 projects in the Proteomics Identifications Database (PRIDE) [17]. The datasets were essentially selected at random, with the constraint that Param-Medic [18] detected variable modifications in 10 of the 20 files.

For search results using Tide, we used `tide-index` 10 times to create 10 different randomly shuffled decoy peptide databases for each PRIDE-20 data set, each of which was paired with the target to yield 10 target-decoy databases. We applied `tide-search` twice to each of the 20 × 10 possible combinations of spectrum file and database: once in a narrow precursor window mode and once in wide precursor window (open search) mode. These settings are described in [6] except, again, the `--concat` option was set to `T` for searches subsequently used by Percolator and by the single-decoy mode Percolator-RESET, and it was to `F` for searches subsequently used by Percolator-RESET’s two-decoy mode.

For search results using MSFragger v3.8, we first converted each of the 20 PRIDE-20 spectrum files from `mgf` to `mzML` format using MSConvert v3.0.23038-42ca1b6. Then we constructed two decoy protein databases for each dataset, one that reversed the target proteins, and another that cyclically permuted the collection of amino acids between any pair of tryptic sites (“K” and “R”) or the terminal protein amino acid. For example, the protein “FGHIJKLMPN” would result in a decoy protein of “JFGHIKPLMN.” For narrow and open searches using MSFragger, we used the default parameter settings except we set the variable modifications and the `fragment mass - tolerance` option according to Param-medic and used no missed cleavages. Searches that were subsequently used by Percolator and by the single-decoy mode Percolator-RESET were conducted against the concatenated target-decoy database twice, once for each of the two constructed decoy databases. Searches that were subsequently used by Percolator-RESET’s two-decoy mode were conducted against the combined database comprised of the target and both decoy databases.

### 2.6 Percolator and Percolator-RESET settings

The `pin` files produced by Tide were processed as described in [6], by filtering for the top 1 PSMs and setting the `enzInt` feature to 0 for all PSMs, because it has been shown that this feature can compromise Percolator’s FDR control [2]. Overall, the features used for training are `deltLCn, deltCn, Xcorr, TailorScore, PepLen, Charge, enzN, enzC, lnNumSP, dM`, and `absdM`, where `Charge` is represented by a one-hot vector indicating the charge state of the precursor (see Supplementary Table 1 for details). No alterations were made to the `pin` file produced by MSFragger search results, and all column features were used. Only the top 1 PSMs were reported by default.

Percolator settings (in Crux) were the same as described in [6, Section 3: Percolator settings]. Subsequent analysis involved first defining the list of considered peptides as done in Percolator-RESET in single-decoy mode (Section 2.3.1) except that we used the Percolator-rescored PSMs rather than the search engine ones. In particular, pairing information was used in the same way it was in Percolator-RESET. Finally, we used TDC to define the list of discovered peptides.

Our Python implementation of Percolator-RESET’s two-decoy mode accepts two distinct formats of the search engines’ output `pin` files. The first, which we used in combination with Tide, takes in two pairs of `pin` files. Each pair corresponds to the target and decoy PSMs obtained by searching the spectra against the target and one of the two decoy databases at a time. In this case we adjust some of the features in the `pin` files so that they are calculated with respect to the combination of the target and two decoy databases rather than separately for each of the two searches. These features include `lnNumSP` (the number of candidate peptides in the combined database for each spectrum), `deltCn` (the difference between the XCorr score of the two top ranked PSMs with respect to the combined database, divided by maximum of the top PSM XCorr score and 1), and `deltLCn` (the difference between the XCorr score of the top scoring PSM and the fifth/last ranked PSM with respect to the combined database, divided by the maximum of the top PSM’s XCorr score and 1). The second input format, which we used in combination with MSFragger, consists of a single `pin` file that contains all the PSMs generated by searching against the concatenated target and two decoy databases. In this case no adjustment are made to the features in the `pin` file.

Finally, because Tide can produce a list of naturally paired target-decoy peptides we could use this information to enable the PSM-and-peptide protocol for defining the considered peptides in both the single-decoy mode (when using Percolator and Percolator-RESET), and the two-decoy mode (Percolator-RESET only). MSFragger, on the other hand, does not provide such pairing, so when defining Percolator’s and Percolator-RESET’s considered peptides in this case we used the PSM-only protocol (`--pair F` in our Python script).

## 3 Results

### 3.1 Percolator-RESET controls the FDR

A given procedure can be said to control the FDR in two complementary ways: theoretically and empirically. A rigorous proof of theoretical FDR control is highly desirable, but even with such a proof, empirical evidence of FDR control is also useful, as a way to check that the assumptions underlying the proof hold in practice.

#### 3.1.1 Theoretical analysis

Consider the abstract setting of the feature-augmented TDC that we are discussing here, where each hypothesis *H*_*i*_ is augmented by additional features or side-information **x**_**i**_. Analogous to the original TDC approach, our meta-procedure RESET assumes that we can compete the original/target score of each hypothesis with a corresponding decoy score so as to assign to each hypothesis a (winning) score *W*_*i*_ and a corresponding target/decoy label *L*_*i*_ = ±1.

In [5] we proved that if the following assumption on the relation between the scores, the labels and the features holds, then RESET controls the FDR in the finite sample setting.

##### Assumption 1.

*Conditional on all the winning scores W, on all the features* **x**, *and on the labels L of the false null hypotheses, the labels of the true null hypotheses are independent identically distributed* ±1 *random variables with P* (*L*_*i*_ = 1) = *P* (*L*_*i*_ = −1) = 1*/*2 *in the single-decoy case, and P* (*L*_*i*_ = 1) = 1 − *P* (*L*_*i*_ = −1) = 1*/*3 *in the two-decoy case*.

Because Percolator-RESET is an adaptation of RESET it inherits its theoretical FDR control. However, this property will only be useful as long as we can translate Assumption 1 to a reasonable equivalent assumption in Percolator-RESET’s context — a task that is somewhat complicated by the fact that, when defining the considered peptides, Percolator-RESET employs either a double competition (PSM-and-peptide) or a single one (PSM-only). In both cases we associate a null hypothesis with each considered peptide, but the nature of these hypotheses changes with the protocol, as explained next.

Consider first the application of Percolator-RESET using PSM-and-peptide in the single-decoy mode. The generated list of considered peptides is made of the winning peptides selected from each target-decoy peptide pair that is associated with at least one PSM. In this case, the null hypothesis we associate with each considered peptide is “the pair’s target peptide is not present in the sample.” If we assume that for each true null hypothesis its competing target and decoy scores are drawn from the same null distribution independently of all other considered peptides, then that assumption suffices to establish that applying TDC to (*W, L*) controls the FDR (in particular, it trivially follows that *P* (*L*_*i*_ = 1) = *P* (*L*_*i*_ = −1) = 1*/*2). Clearly, the validity of this assumption depends on how the decoys are generated, and at best it is a reasonable approximation (we will revisit this point in the Discussion). Regardless, this assumption falls short in addressing RESET’s use of the additional features.

Recall that in our case the (additional) features **x**_*i*_ are inherited from the highest scoring PSM attached to the target-decoy pair. Therefore, our strengthened assumption is

##### Assumption 2.

*For each true null hypothesis, its competing target and decoy scores together with their additional features are drawn from the same null distribution independently of all other considered peptides*.

While plausible, the validity of the last assumption now depends not only on how the decoys are generated, but also on which features are used. Poorly selected features — as an extreme example consider a feature that identifies the peptide as a target or a decoy — can clearly violate our assumption. Regardless, if our last assumption holds then it is easy to see that so does Assumption 1 and therefore Percolator-RESET controls the FDR among its list of reported peptides in this case.

Adjusting the last discussion to the application of Percolator-RESET using PSM-and-peptide in the two-decoy mode is straightforward. Indeed, in this case the generated list of considered peptides is made of the winning peptides selected from each target-decoy-decoy matched peptide triplet that is associated with at least one PSM. Therefore, the null hypothesis is modified to “the triplet’s target peptide is not present in the sample.” Similarly, our assumption needs to be modified to

##### Assumption 3.

*For each true null hypothesis, its competing target and one / two decoy scores (according to single- or two-decoy mode) together with their additional features are drawn from the same null distribution independently of all other considered peptides*.

Again, if this latter assumption holds then so does Assumption 1 (in particular it trivially implies that *P* (*L*_*i*_ = 1) = 1 − *P* (*L*_*i*_ = −1) = 1*/*3), thus establishing rigorous FDR control in this case.

Consider next the application of Percolator-RESET using PSM-only (i.e., without any pairing information) in the single-decoy mode. The list of considered peptides is now made of all the peptides that are associated with at least one PSM. For the purpose of using our theorem we need to modify the null hypothesis we associate with each considered peptide to “the PSM representing the peptide is incorrect.” However, Assumption 2 remains the same. Indeed, all that changes is what we mean by a true null hypothesis (as explained), and by “competing target and decoy scores”: whereas before we referred to the paired peptide scores, here we refer to the best target and decoy peptides matching the considered spectrum.

If the last assumption is satisfied then again we can invoke Theorem 1 of [5] to ensure us that Percolator-RESET controls the FDR when using PSM-only. However, there is a subtle difference here compared to when using PSM-and-peptide. Specifically, the null assumption changed, so whereas a true null hypothesis for PSM-and-peptide corresponds to a target peptide that is not present in the sample, for PSM-only it corresponds to an incorrect PSM representing the peptide. Therefore, the false discoveries in this case are peptides whose representing PSM is incorrect, but this does not preclude that the peptide itself is in the sample. Fortunately, this only makes Percolator-RESET conservative when using PSM-only because the procedure is more strict in the type of error it controls among its reported list of peptides. In particular, it means that under our last assumption Percolator-RESET still rigorously controls the FDR.

Finally, adjusting the above analysis to the application of Percolator-RESET using PSM-only in the two-decoy mode is again straightforward. The null hypotheses remain unchanged, and again we generalize Assumption 2 to Assumption 3, which remains the same as in the two-decoy mode with PSM-and-peptide but its semantics are modified analogously to the single-decoy case.

#### 3.1.2 Entrapment experiments

We supplemented our theoretical analysis of Percolator-RESET’s FDR control with a series of entrapment experiments described in Section 2.4. Figures 1A-B and 2A-B summarize the results of these experiments by comparing the selected FDR threshold with the empirical FDR. Notably, Percolator-RESET’s empirical FDR stays below or at most just above the black diagonal line which coincides with the prescribed threshold. This observation holds for all proportions of the in-sample peptides (these varying proportions correspond to varying the signal-to-noise in our searches), regardless of whether Tide was employed in open or narrow search mode, whether Percolator-RESET was applied in single-decoy or in two-decoy mode, and whether it was applied with or without pairing information.

**Figure 1:**
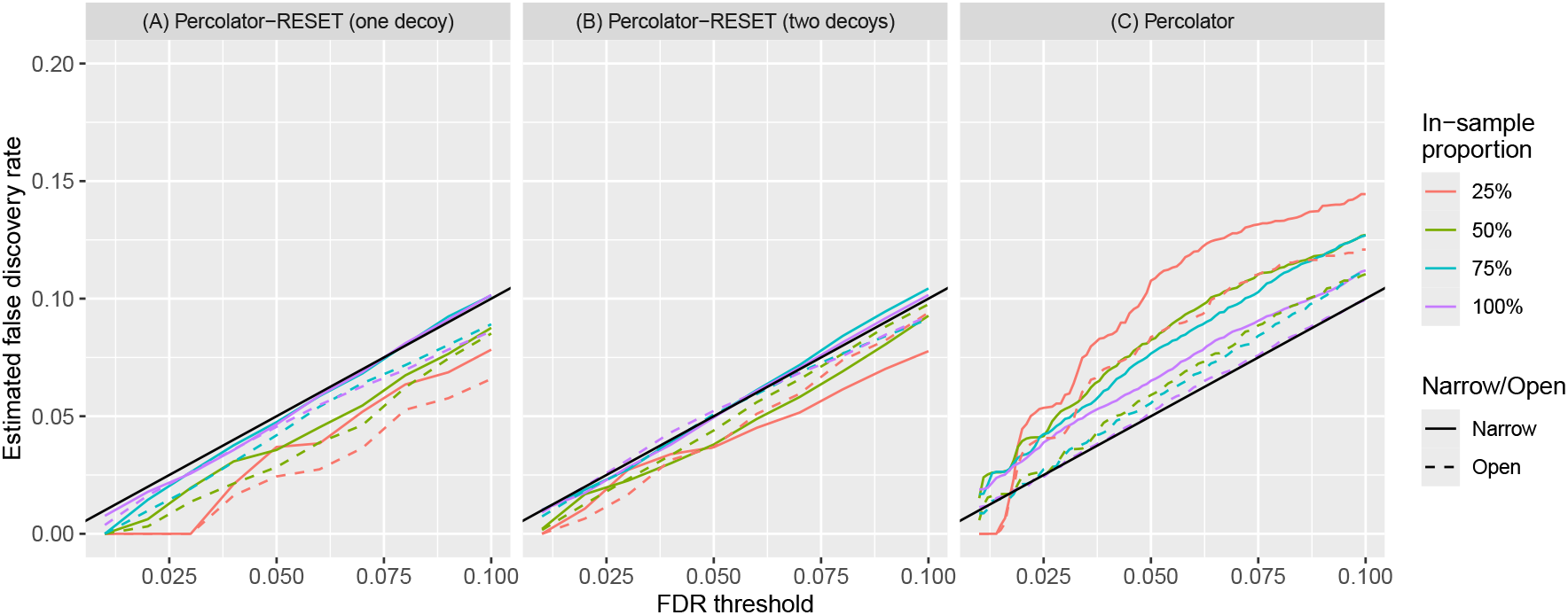
Empirical FDRs of Percolator Algorithms. The empirical FDR as estimated from our entrapment experiments on the ISB18 dataset using (A) single-decoy Percolator-RESET, (B) two-decoy Percolator-RESET, (C) Percolator. The FDP is estimated at a range of FDR thresholds ([0.01, 0.1]), and its average over 100 randomly generated decoys (and over 2 randomly generated training decoy sets for (A) and (B)) is the empirical FDR. Each curve represents the empirical FDR using a target database constructed with the specified proportion of the in-sample database, in either narrow- or open-search mode. All peptide-level analyses were using the PSM-and-peptide protocol utilizing target-decoy pairing/matching information.

**Figure 2:**
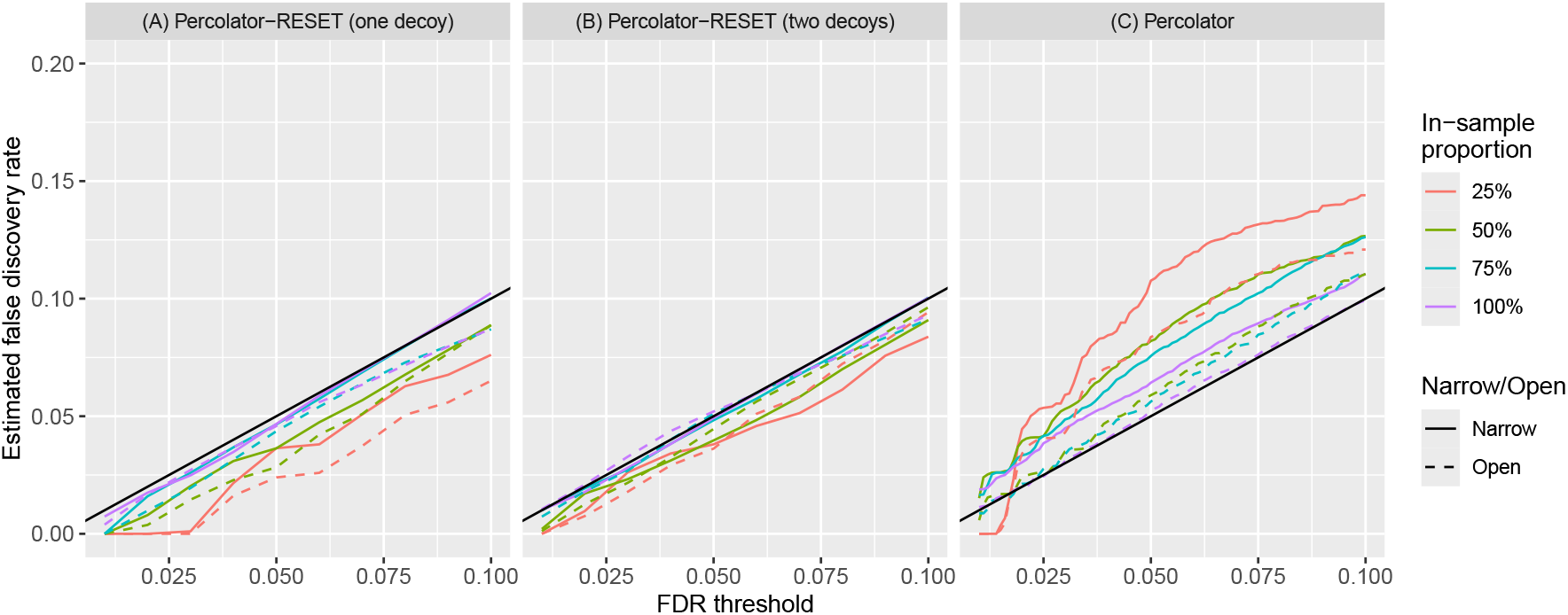
Empirical FDRs of Percolator Algorithms without target-decoy pairing/matching information. The empirical FDR as estimated from our entrapment experiments on the ISB18 dataset using (A) single-decoy Percolator-RESET, (B) two-decoy Percolator-RESET and (C) Percolator using the PSM-only protocol which does not require any pairing information. The FDP is estimated at a range of FDR thresholds ([0.01, 0.1]), and its average over 100 randomly generated decoys (and over 2 randomly generated training decoy sets for (A) and (B)) is the empirical FDR. Each curve represents the empirical FDR using a target database constructed with the specified proportion of the in-sample database, in either narrow- or open-search mode.

More specifically, as a percentage, the worst violation of Percolator-RESET was recorded when it was applied using PSM-and-peptide in the two-decoy mode to open search results generated by Tide using a database that contained 100% of the ISB18 peptides, where at an FDR threshold of 0.04, the empirical FDR was 0.0431. Given that the empirical FDR was obtained by averaging the FDP over 100 decoys, this 8% violation does not seem that significant. Moreover, keep in mind that even if we were able to average the FDP over a much larger number of decoys, we still do not average out all the variability in the experiment: the MS/MS runs themselves introduce a random effect that cannot be eliminated just by averaging over multiple drawn decoys. Indeed, accounting for that latter effect would have required repeating these MS/MS experiments many times.

We also reanalyzed Percolator’s behavior using this entrapment setup. The results in Figure 1C and Figure 2C, done here with 100 decoy databases, reinforce our previously reported observation (based on only 20 decoy databases), that Percolator struggles to control the FDR in these entrapment experiments [7]. The worst empirical FDRs across the two figures at thresholds 0.01, 0.05 and 0.1 were 0.0189, 0.1077 and 0.1445, respectively. As noted previously, we believe that Percolator’s failure to control the FDR stems from the presence of multiple spectra generated from the same peptide species. Consequently, these essentially identical spectra may appear in both the training and test sets in Percolator’s cross-validation scheme causing the model to overfit and violate FDR control.

### 3.2 Percolator-RESET and Percolator report a similar number of discoveries

Percolator can fail to control the FDR, and therefore its results should be taken with a grain of salt. However, particularly in light of Percolator’s popularity, it is instructive to compare its apparent statistical power with Percolator-RESET’s. Specifically, we examined the ratio of the number of Percolator-RESET to Percolator discoveries when both tools were applied in a variety of settings to multiple search results. These results were obtained by applying Tide and MSFragger, in both narrow and open search modes, to our PRIDE-20 collection of 20 publicly available MS/MS datasets, and using multiple decoy databases (Section 2.5).

In general we found that, as we vary the FDR threshold between 1–5%, there is little that separates the two tools power-wise: the Percolator-RESET to Percolator discoveries ratio typically hovers around 1. Figures 3-4 summarize those findings by displaying the quartiles of the averaged ratios of discovery numbers, where the quartiles are taken with respect to the 20 PRIDE-20 datasets and the average with respect to varying the decoy databases.

**Figure 3:**
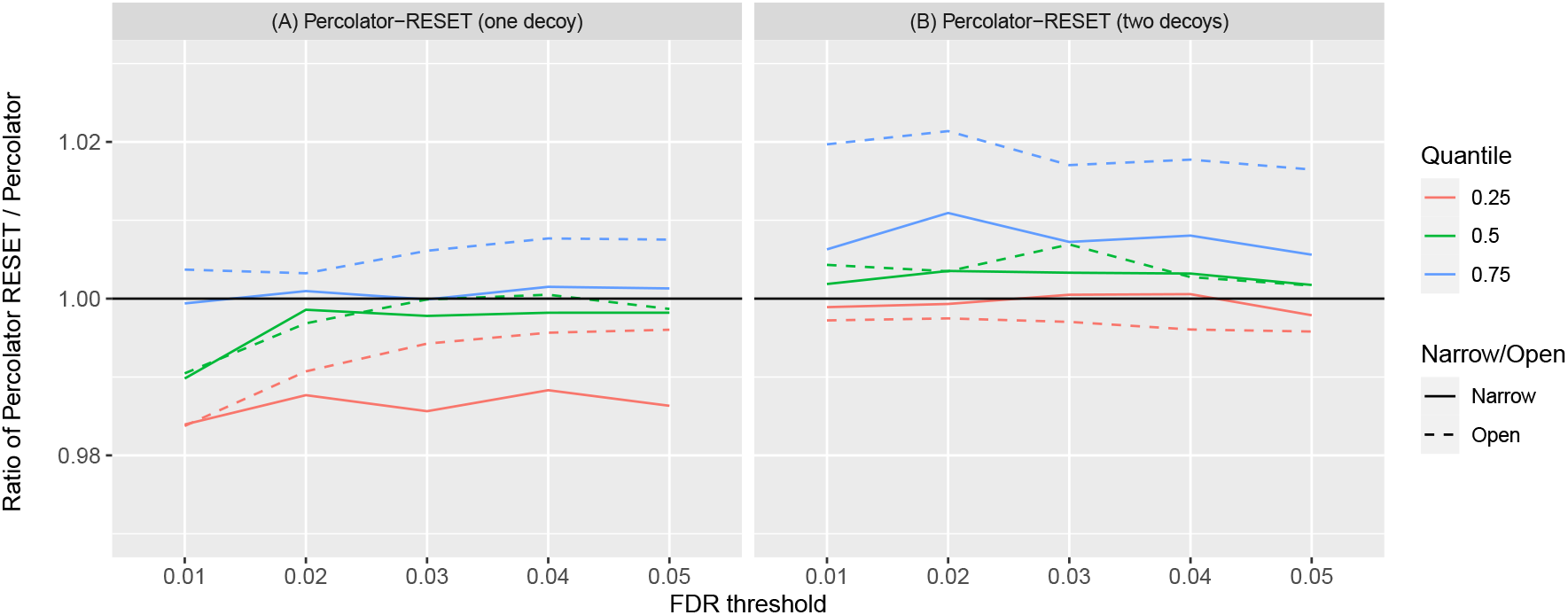
Percolator-RESET procedures discover similar number of peptides compared to Percolator using Tide. The left panel displays the quartiles, with respect to the 20 PRIDE-20 datasets, of the mean ratio of the single-decoy Percolator-RESET discovered peptides to Percolator discovered peptides. The right panel similarly covers the mean ratio of the two-decoy Percolator-RESET discovered peptides to Percolator discovered peptides. Searches were conducted using Tide search. For the left panel, the mean ratio is taken with respect to each dataset’s 10 decoys. For the right panel, because this version of Percolator-RESET considers two decoy databases, we calculated the number of discoveries 10 times, each using a unique pair from the 10 available decoy databases, while ensuring that each decoy database was used exactly twice. Subsequently, the discovery ratio in that case is computed as the number of discoveries of the two-decoy Percolator-RESET to the average number of discoveries of Percolator using the two decoy databases separately.

**Figure 4:**
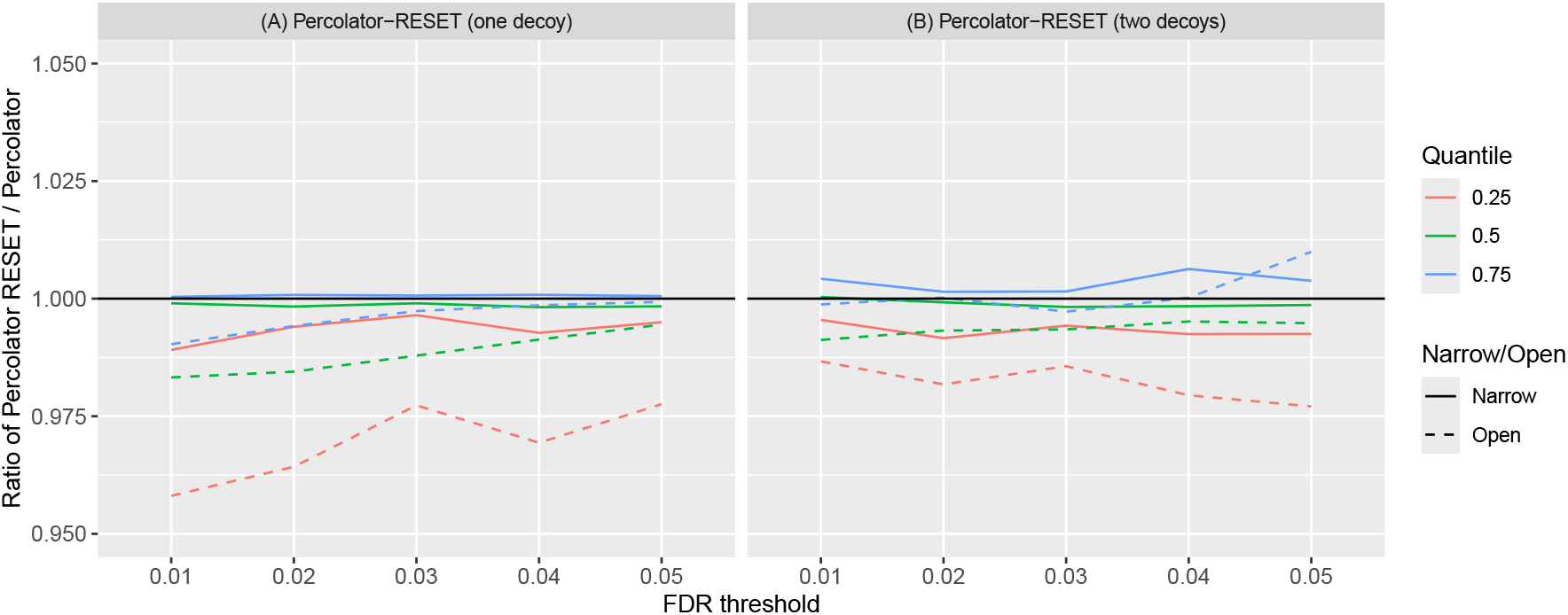
Percolator-RESET discovers similar number of peptides compared to Percolator without target-decoy pairs in MSFragger. (A) the quartiles, with respect to the 20 PRIDE-20 datasets, of the mean ratio of the single-decoy Percolator-RESET discovered peptides to Percolator discovered peptides. (B) quartiles of the mean ratio of the two-decoy Percolator-RESET discovered peptides to Percolator discovered peptides. Searches were conducted using MSFragger and the lists of considered peptides were constructed without any pairing information. In (A) we used 10 applications of Percolator-RESET (single decoy) while randomly varying the training/estimating decoy-splitting, 5 times for each of the 2 decoy databases constructed. We applied Percolator twice, once for each decoy database. Then for each decoy database, we took the ratio between each of the 5 Percolator-RESET runs and Percolator, and then averaged the results. In (B) we used 10 applications of Percolator RESET’s two-decoy mode, using both decoy databases in each application, again varying the training/estimating decoys. Since Percolator only uses one decoy-database at a time, we averaged the number of discoveries of the two Percolator runs. Then for each of the 10 applications, we took the ratio of the two-decoy Percolator-RESET and the averaged number of discoveries from Percolator, and then averaged these results.

Looking more closely, Figure 3 compares the tools when both are applied with target-decoy pairing information (using PSM-and-peptide) to Tide’s search results. For panel (A) each average ratio was obtained by averaging the ratios observed from 10 applications of Percolator and of Percolator-RESET (in single decoy mode). The median average ratio (green) for both narrow and open searches is between 0.99–1. Percolator-RESET’s weakest performance here is at the 1% FDR threshold, where its median power across the PRIDE-20 dataset is 99% of Percolator’s in both narrow and open search mode. At higher FDR thresholds, this difference in number of discoveries becomes even more marginal, with the median average ratio at the 5% FDR threshold at 99.8% and 99.9% in narrow mode and open mode, respectively. Moreover, in panel (B) the two-decoy mode of Percolator-RESET marginally edges out Percolator, with median average ratios of 100.2% and 100.4% at the 1% FDR threshold, and a median greater than 100% for all remaining FDR thresholds. The slight improvement is perhaps not surprising, given that the algorithm uses an additional decoy database.

Note that in panel (B) we used Percolator-RESET’s two-decoy mode, so the average ratio of discoveries was defined differently. Specifically, each of the 10 applications of Percolator-RESET used a unique pair of the 10 available decoys so that overall each decoy was used exactly twice. Of course, Percolator only uses a single decoy, so the ratio here was defined as the number of Percolator-RESET discoveries divided by the average number of corresponding Percolator discoveries using the two decoy databases separately. Averaging those 10 ratios over the 10 decoys pairs defined the average ratio.

Figure 4 offers a similar comparison of the tools but applied without any pairing information (using PSM-only) to process MSFragger’s searches. In this scenario Percolator-RESET is relatively slightly weaker than when using the paired information and Tide searches, but the power loss relative to Percolator is still essentially marginal. Indeed, at the 1% FDR level, the mean ratio of Percolator-RESET (single decoy) to Percolator discoveries has a median of 99.9% and 98.5% using narrow and open search modes respectively (A), and in the two-decoy mode the medians are 100% and 99.1% (B).

Given that our MSFragger searches used only two decoys for each dataset we had to revise the definition of average ratio once again. In (A), using each one of the two decoys at a time we applied Percolator-RESET five times, randomly varying the training/estimating decoy-split each time. We next averaged those number of discoveries across the five applications and divided it by the corresponding number of Percolator discoveries using that decoy. Finally we averaged those two ratios to define the average ratio. In (B) Percolator-RESET was engaged in the two-decoy mode, so to define the average ratio we first apply it 10 times to both decoys while randomly varying the training/estimating decoy-split each time. In each case, we divided the number of discoveries by the corresponding number of Percolator discoveries averaged over the two decoys. The average of those 10 ratios over the 10 random decoy splits defined the average ratio.

### 3.3 Double competition provides only a marginal boost for Percolator-RESET

We previously reported that when using TDC for peptide-level analysis, the double competition protocol, PSM-and-peptide, tends to deliver more discoveries than the single competition, PSM-only, protocol. Moreover, we demonstrated that the difference can at times be substantial [16]. Thus, it is interesting to note that while the use of PSM-and-peptide in defining Percolator-RESET’s list of considered peptides is still advantageous, it delivers only a marginal power boost over using PSM-only.

This effect is clearly visible in Figure 5, which looks at the quartiles (with respect to the PRIDE-20 datasets) of the average ratio of Percolator-RESET discoveries with pairing (PSM-and-peptide) to without pairing (PSM-only). In both cases, Percolator-RESET is applied to the same Tide open and narrow searches that we previously analyzed when comparing to Percolator. The average ratio in the single-decoy mode (A) is the average of the corresponding 10 discovery ratios, one for each of the 10 decoy databases we used. For the two-decoy mode (B) we average the ratio of with-to-without pairing discoveries in 10 applications of Percolator-RESET, each applied to a unique pair of decoys, where each of the 10 available decoys appears in exactly two such pairs.

**Figure 5:**
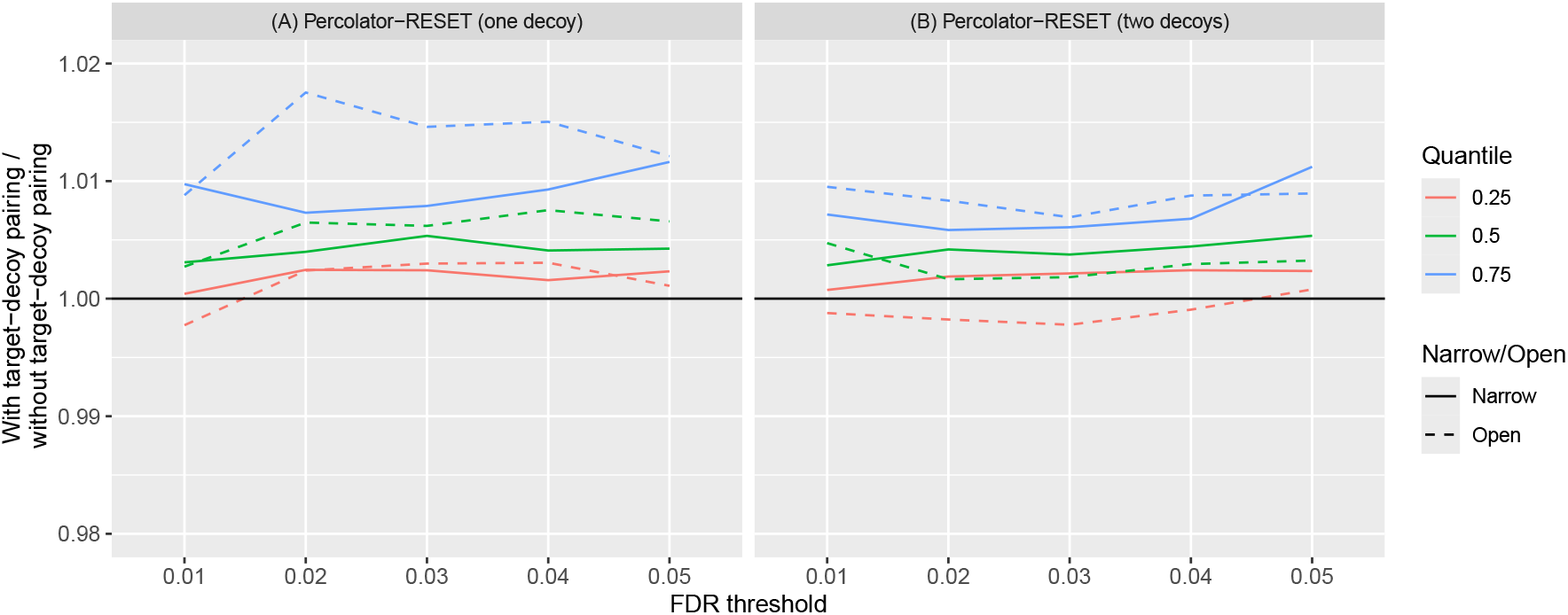
Percolator-RESET gets a marginal power boost when using target-decoy pairing information. Both panels display the quartiles, with respect to the 20 PRIDE-20 datasets, of the mean ratio of Percolator-RESET discovered peptides with vs without the second round of peptide-level competition. Searches were conducted using Tide search. (A) Percolator-RESET was applied in single-decoy mode and the mean ratio is taken with respect to each dataset’s 10 decoys. (B) Percolator-RESET was applied in two-decoy mode so we calculated the number of discoveries 10 times, each using a unique pair from the 10 available decoy databases, while ensuring that each decoy database was used exactly twice.

The marginal power gain in pairing here could possibly be attributed to the improved ranking of Percolator-RESET. Indeed, the second-level competition helps in eliminating some of the relatively high-scoring decoys that still score lower than their paired targets. It could be that Percolator-RESET achieves the same goal by simply lowering the rank of such decoys rather than eliminating them altogether. Regardless, the reassuring message is that when target-decoy pairing is unavailable, as is typically the case, the users should expect only a marginal decrease in the number of Percolator-RESET discoveries.

### 3.4 Percolator-RESET’s two-decoy mode offers reduced variability

Percolator-RESET has an intrinsic variability due to the randomized splitting of the decoy set into training and estimating decoys: unless its random number generator seed is set to a fixed value, the procedure typically produces somewhat different results in different applications to the same dataset. Although fixing the seed eliminates the variability, we do not recommend doing so if for no other reason than the theoretical guarantees of the FDR control would no longer apply in this case. However, the two-decoy mode offers a viable alternative to reducing the variability.

Indeed, in this mode each target has to beat both competing decoys rather than just one. Because the sets of training and estimating decoys are roughly equal-sized, both sets are therefore typically much larger in this mode. While these larger training and estimating sets come at a cost of a reduced number of target wins, keep in mind that these lost targets only win one of their two decoy competitions; hence, they are unlikely to be high scoring enough to eventually make it to the reported discoveries list. At the same time, those larger sets of training and estimating decoys should allow us to better train our model, as well as to have a more stable final procedure when defining the discovery list using `SSS+`. This prediction is borne out in a couple of experiments that are described next.

We began the first experiment by finding, for each PRIDE-20 spectrum file, the sample variance in the number of discoveries reported in 10 applications of the single-decoy version of Percolator-RESET, each using a different decoy database. We then compared each of those 20 sample variances with corresponding sample variances of 10 applications of the two-decoy Percolator-RESET, each using a distinct pair of decoy databases (each of the 10 available decoy databases was used in exactly two applications). Specifically, for each PRIDE-20 spectrum file we calculated the ratio of the two-decoy Percolator-RESET to the single-decoy Percolator-RESET sample variances. Figure 6A shows the quartiles (over the 20 datasets) of these ratios in logarithmic scale. At the 1% FDR threshold the two-decoy mode typically exhibits substantially less variability than the single-decoy mode, yielding a median reduction of 29.8% in the narrow-search and 63.4% in the open search.

**Figure 6:**
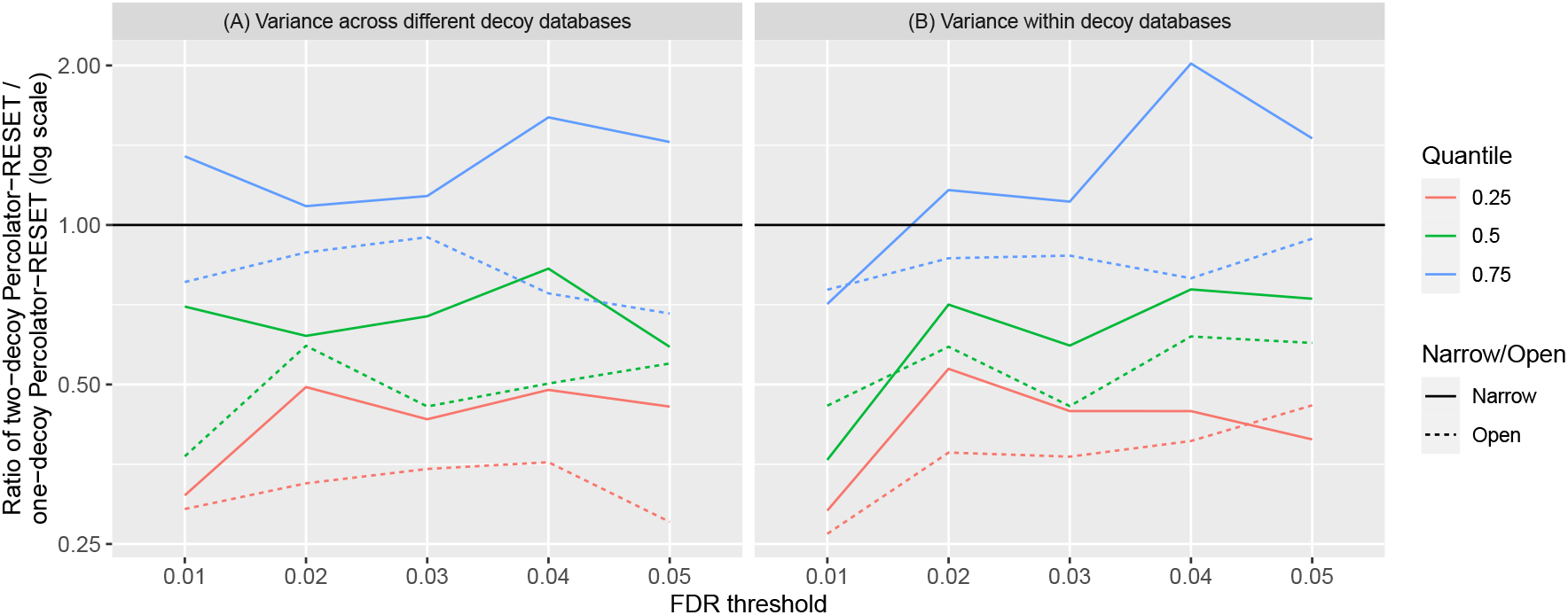
Comparing the variabilities of Percolator-RESET in single- and two-decoy modes. Both panels display the quartiles, taken over the 20 PRIDE-20 datasets, of the log_2_ ratios of variances in the number of discoveries from two-decoy Percolator-RESET (numerator) to one-decoy Percolator-RESET (denominator). Each PRIDE-20 dataset had 10 decoy databases constructed. In (A), we calculated the sample variance for the two-decoy mode using 10 applications, with each application using a unique pair selected from the 10 available decoy databases. The sample variance for the single-decoy mode was calculated using 10 applications, each using a single decoy database from the 10 available decoy databases. The ratios are calculated as the sample variance of the two-decoy mode to the single-decoy mode. In (B), for each PRIDE-20 dataset, we preselected two decoy databases from the 10 available. We ran the two-decoy mode 20 times using the preselected pair of decoy databases, while randomly varying the training/estimating decoy-splitting, and calculated the sample variance in the reported number of discoveries. We ran the single-decoy Percolator-RESET 10 times for each of the two decoy databases (a total of 20), and calculated the pooled sample variance: the average of the sample variance of the number of discoveries for each of the two sets of 10 applications. The ratios are then calculated as the sample variance of the two-decoy mode to the pooled sample variance of the single-decoy mode.

In the second experiment, we focused on the variation in the number of discoveries due to the random splitting of the decoys into training and estimating decoys. For each PRIDE-20 spectrum file we first pre-selected two decoy databases arbitrarily. Next, we applied the two-decoy Percolator-RESET 20 times, each using a different random split between the training and estimating decoys, and we calculated the sample variance in the number of discoveries over these 20 runs. For the single-decoy version, we applied it 10 times using the first decoy database and 10 times using the second decoy database, randomly splitting the decoys in all those applications. Subsequently, we calculated the pooled sample variance in the number of discoveries reported by the single decoy Percolator-RESET, i.e., the average of the two sample variances calculated with respect to the 10 runs using each of the two databases. Finally, we calculated the ratio of the two-decoy mode sample variance to the single-decoy mode pooled sample variance for each PRIDE-20 spectrum file. Figure 6B plots the quartiles of these ratios in logarithmic scale while varying the FDR threshold. Again, we see that Percolator-RESET’s two-decoy mode typically exhibits substantially lower variability than the single-decoy mode, with a median reduction of 64.0% and 54.4% at the 1% FDR threshold when using a narrow and open search, respectively.

## 4 Discussion

The Percolator software takes as input tandem mass spectrometry search results and performs two distinct tasks: first, it uses machine learning to automatically re-rank the given set of PSMs, and second, it uses TDC to estimate the FDR in the resulting list. Our previous analysis [7], as well as our analysis in Section 3.1, demonstrate that Percolator can link these two tasks in a problematic fashion, leading to underestimation of the FDR in the context of peptide detection.

One specific problem we previously highlighted with Percolator arises when multiple PSMs are generated by the same analyte. In this case, essentially the same PSM can end up in both Percolator’s training and testing folds, thereby negating the intended effect of its cross-validation scheme. Moreover, as we pointed out, this problem is exacerbated when, as is often the case, multiple runs are combined. Notably, this problem is not specific to Percolator: any post-processor that uses cross-validation that randomly splits all the PSMs into their folds can inadvertently learn to distinguish between target and decoy PSMs rather than between correct and incorrect ones.

Admittedly, the multiplicity problem can be addressed if Percolator would switch from PSM-level training to peptide-level training, which is also the level Percolator-RESET works on. However, this modification still does not address the fact that the way Percolator combines its three folds is, as of now, not supported by rigorous mathematical theory. As such, a peptide-level variant of Percolator might still possibly fail to control the FDR in some scenarios. In contrast, Percolator-RESET provably controls the FDR under mild assumptions analogous to those that TDC relies on. Moreover, Percolator-RESET’s two-decoy mode can be adjusted so that it can train its SVM at the PSM level while still rigorously controlling the FDR. This is something that neither Percolator nor Percolator-RESET’s single-decoy mode can accomplish. Indeed, the same multiplicity problem makes it impossible to rigorously control the FDR at the PSM-level even when using TDC [10, 16]. We plan on using the adjusted two-decoy mode to investigate the difference between peptide- and PSM-level training in this case.

Theoretical guarantees aside, we show that both the single- and two-decoy modes of Percolator-RESET appear to effectively control the FDR in the same entrapment experiments that Percolator failed to do so. Importantly, in spite of its tighter FDR control, the power of the single-decoy mode of Percolator-RESET is only marginally lower than Percolator’s. Moreover, its two-decoy mode is arguably marginally more powerful than Percolator, while exhibiting significantly less variability than the single-decoy version in the number of discoveries it reports. Regarding the variability, it is worth noting that Percolator-RESET exhibits some while Percolator does not, but that is only because Percolator effectively hides its inherent variability by using a fixed seed so that its stochastic behavior (e.g., in the definition of its cross-validation folds) is masked. The user can specify a seed to Percolator-RESET if they want to similarly hide its variability.

By splitting the considered peptide decoys into training and estimating ones, Percolator-RESET circumvents Percolator’s need for a cross-validation scheme for controlling the FDR (cross-validation is still used for selecting the hyperparameters). Aside from allowing for provable FDR control, dispensing with the cross-validation is also potentially more computationally efficient. Such efficiency could be particularly useful if we want to take advantage of RESET’s flexibility to be used in conjunction with other semi-supervised machine learning models that rely on a negative and an unlabelled set. It would be interesting to compare the impact of different such models on the performance of RESET.

Finally, a word of caution: Percolator-RESET’s rigorous FDR control hinges on the validity of Assumption 3. There are two components to this assumption. The first is the usual one that any variant of TDC depends on, namely, that the decoys are constructed in such a way that for a true null the target and decoy scores are drawn from the same null distribution. In practice this could be very difficult to achieve in some cases. For example, the usual construction of decoys ignores the possible impact of “neighbors”, i.e., peptides that might generate very similar theoretical spectra: a target peptide that is missing from the sample might be mistaken for a close neighbor that is present, so even though the missing peptide represents a true null, it is much more likely to win than its decoy is. Of course this is a potential problem with all variants of TDC and one which in practice is glossed over as being typically negligible. Notably, Percolator-RESET follows CONGA’s footsteps in implementing dynamic-level competition, which alleviates the problem when variable modifications are considered. The second component of Assumption 3 is that the features should not help us distinguish between targets and decoys for true nulls. Indeed, features such as those would compromise our approach, similar to how they can foil Percolator’s FDR control [2, 8]. An interesting topic for future research is to develop tools that can detect such problematic features, particularly because their problematic nature is often only obvious in hindsight.

## A Supplement

### A.1 Features used by Percolator-RESET

### A.2 Algorithms

#### Algorithm 1

**RESET: a meta procedure for feature-augmented TDC**

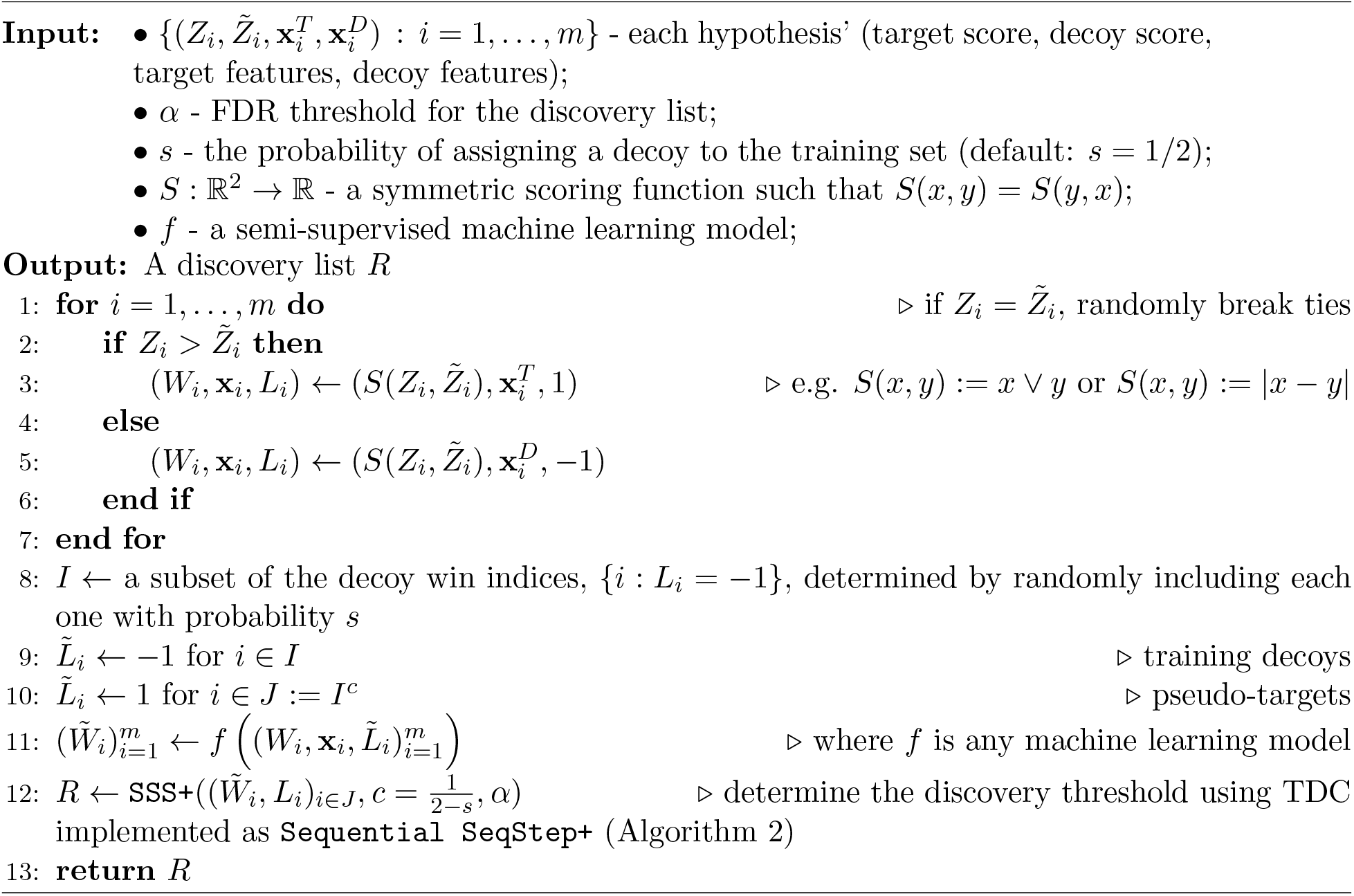

#### Algorithm 2

**Sequential SeqStep / SeqStep+** (adopted from Selective Sequential Step+ of [1])

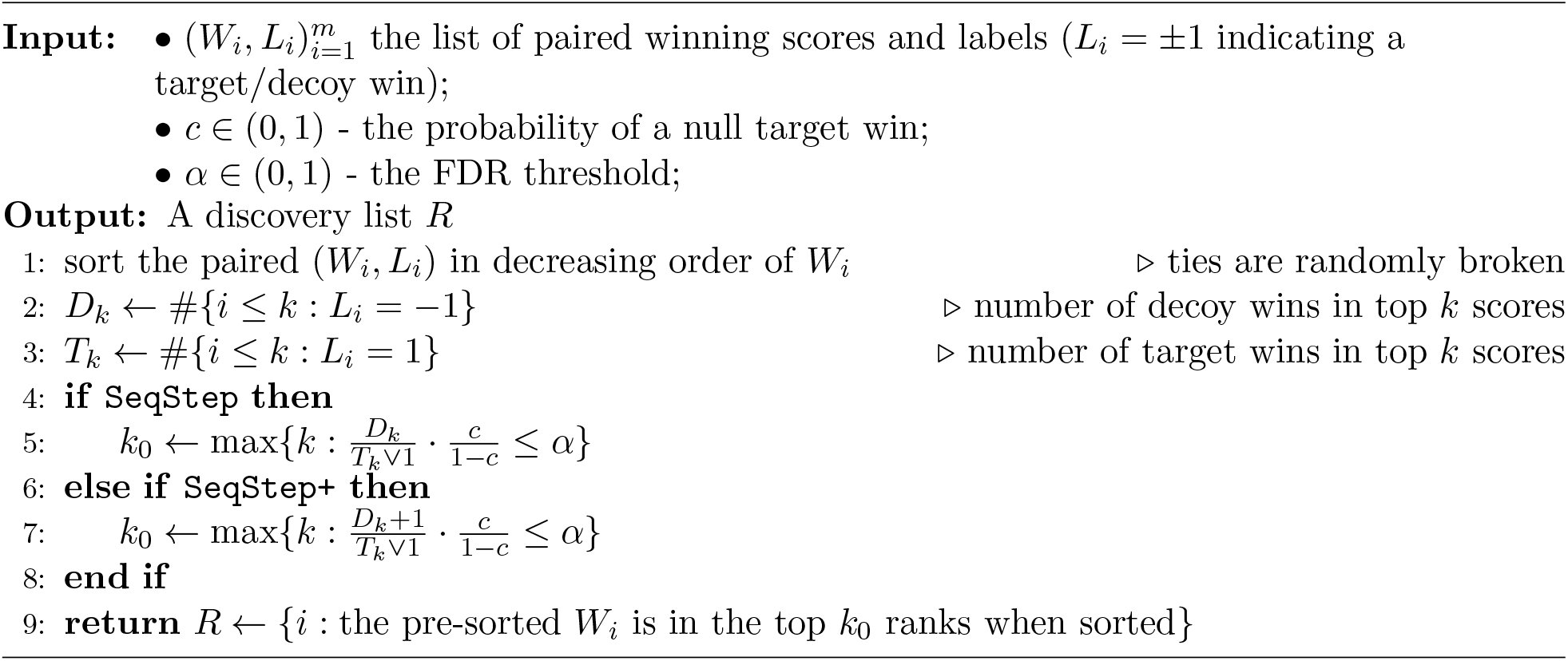

#### Algorithm 3a

**Define the considered peptides and their scores (using 2 decoys)**

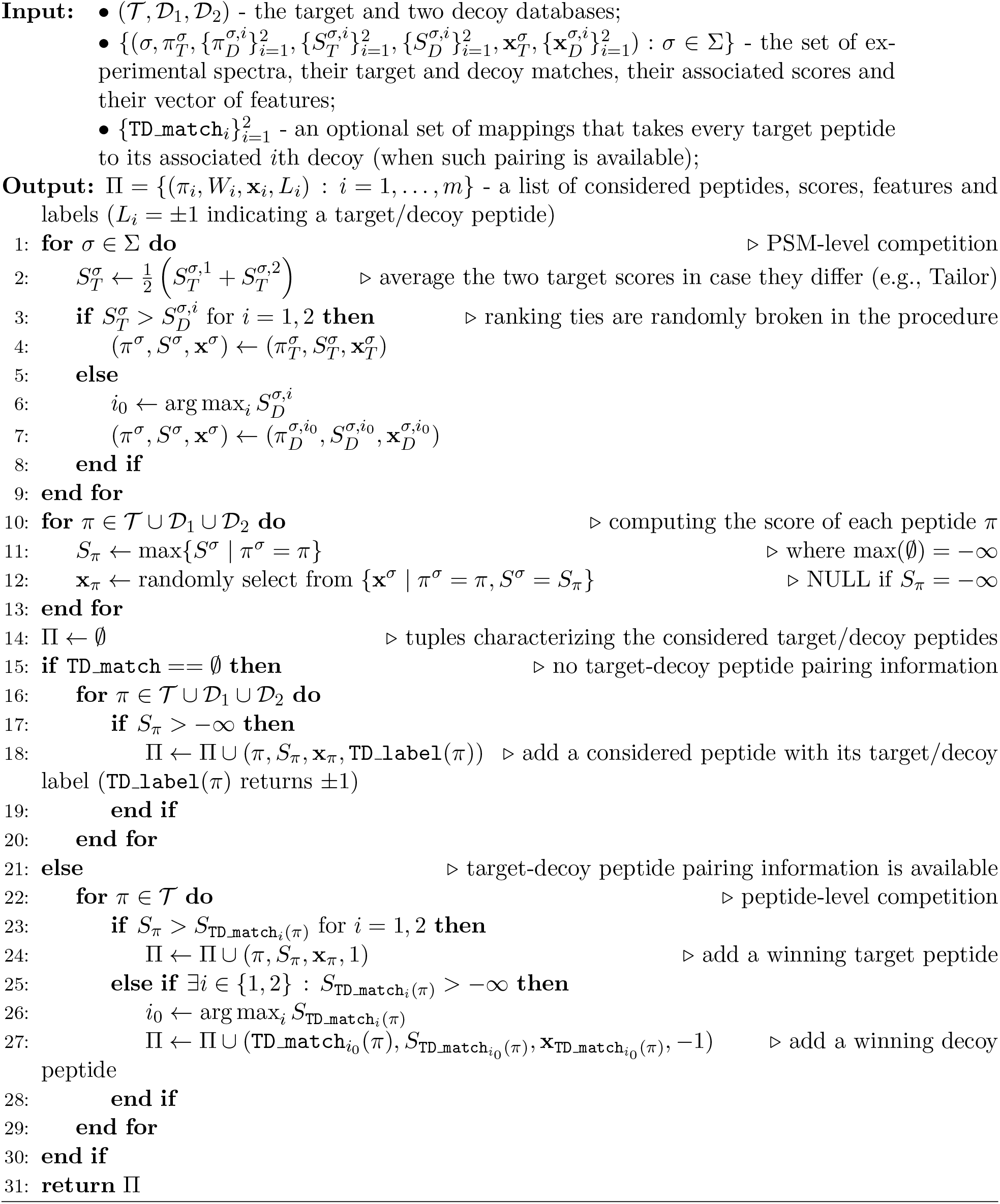

#### Algorithm 3b

**Define the considered representative peptides and their scores (using 2 decoys w/variable modifications)**

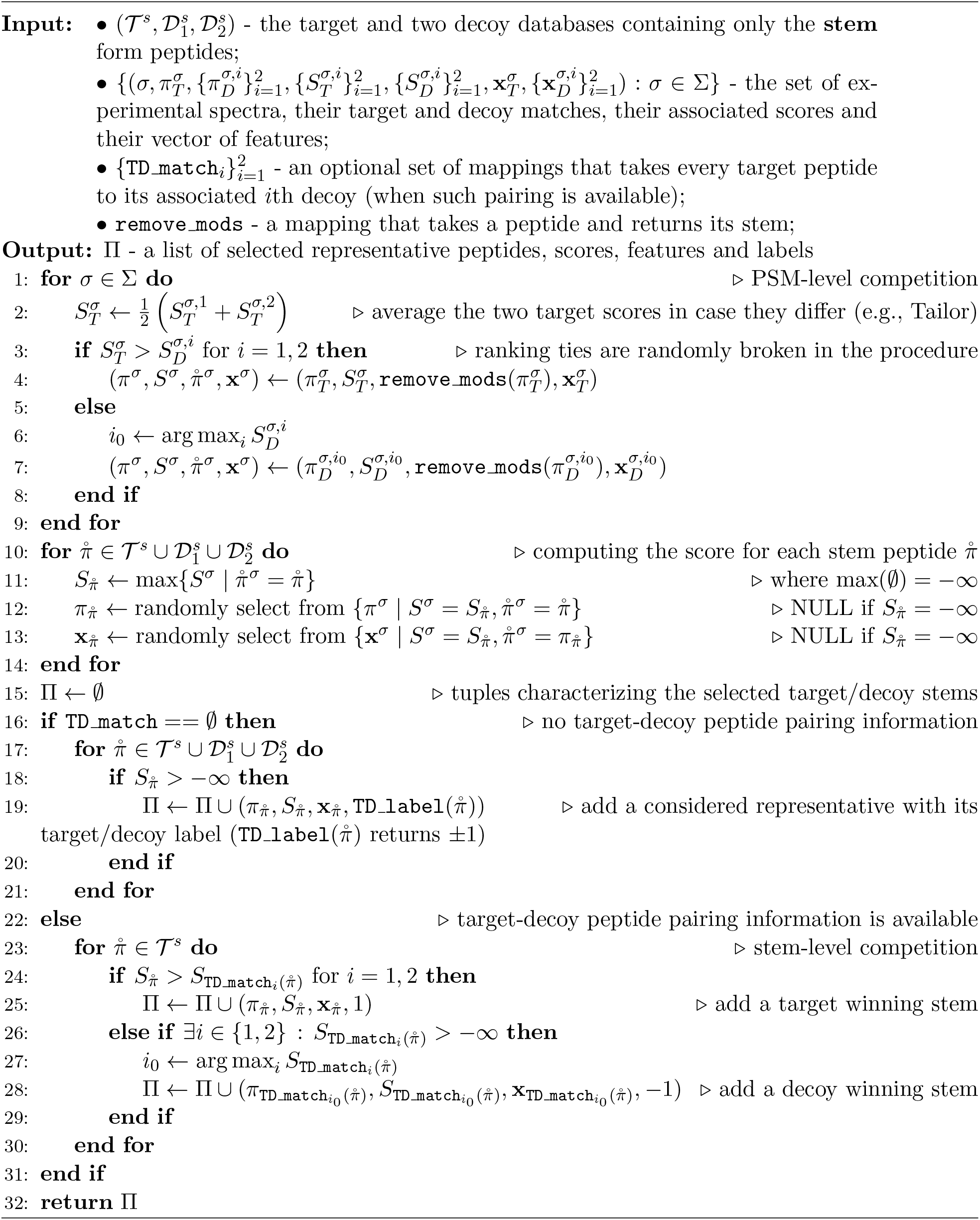

#### Algorithm 4

**Percolator-RESET’s rescoring engine**

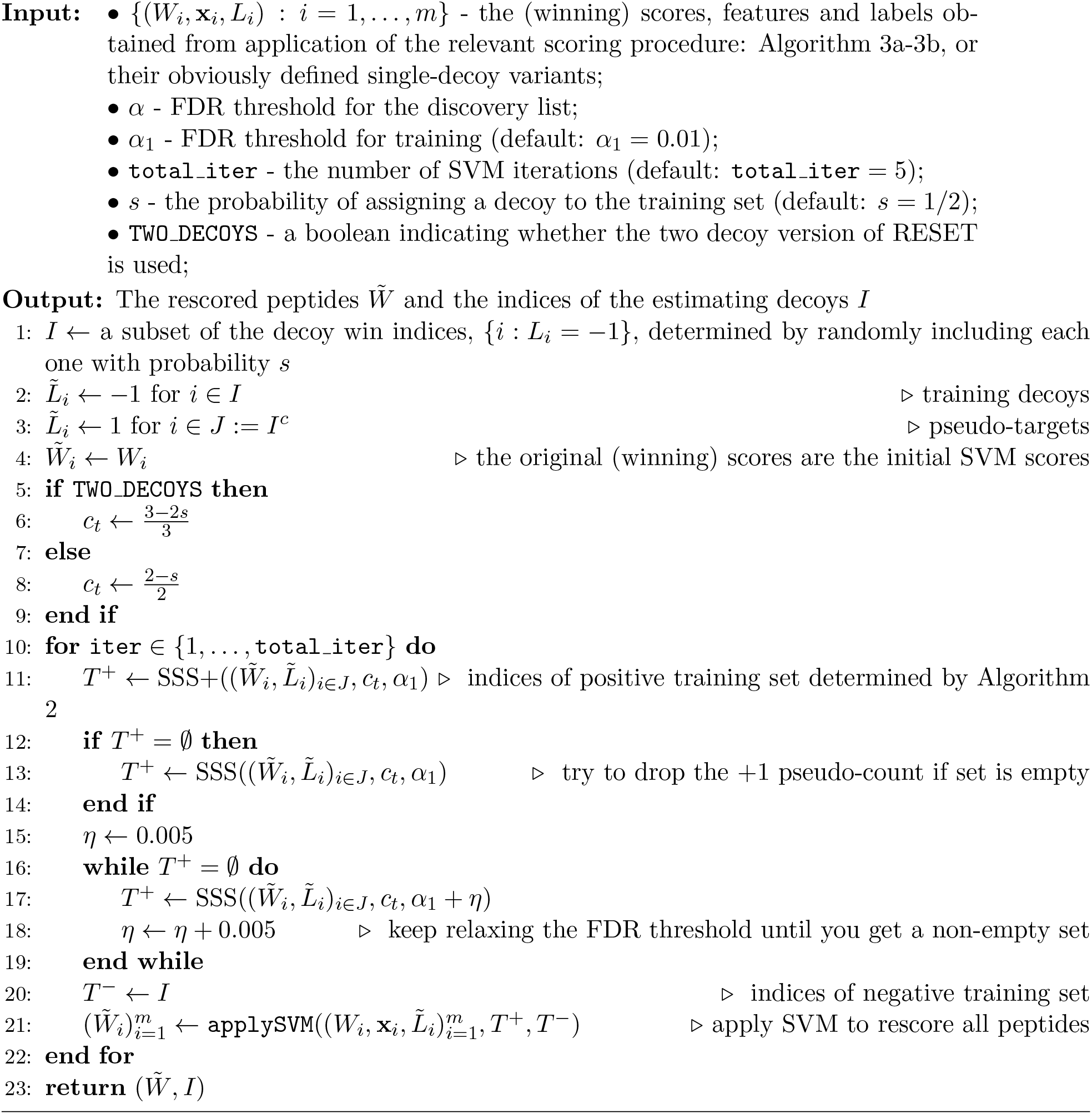

#### Algorithm 5

**Precursor mass clustering (`precursor clustering`)**

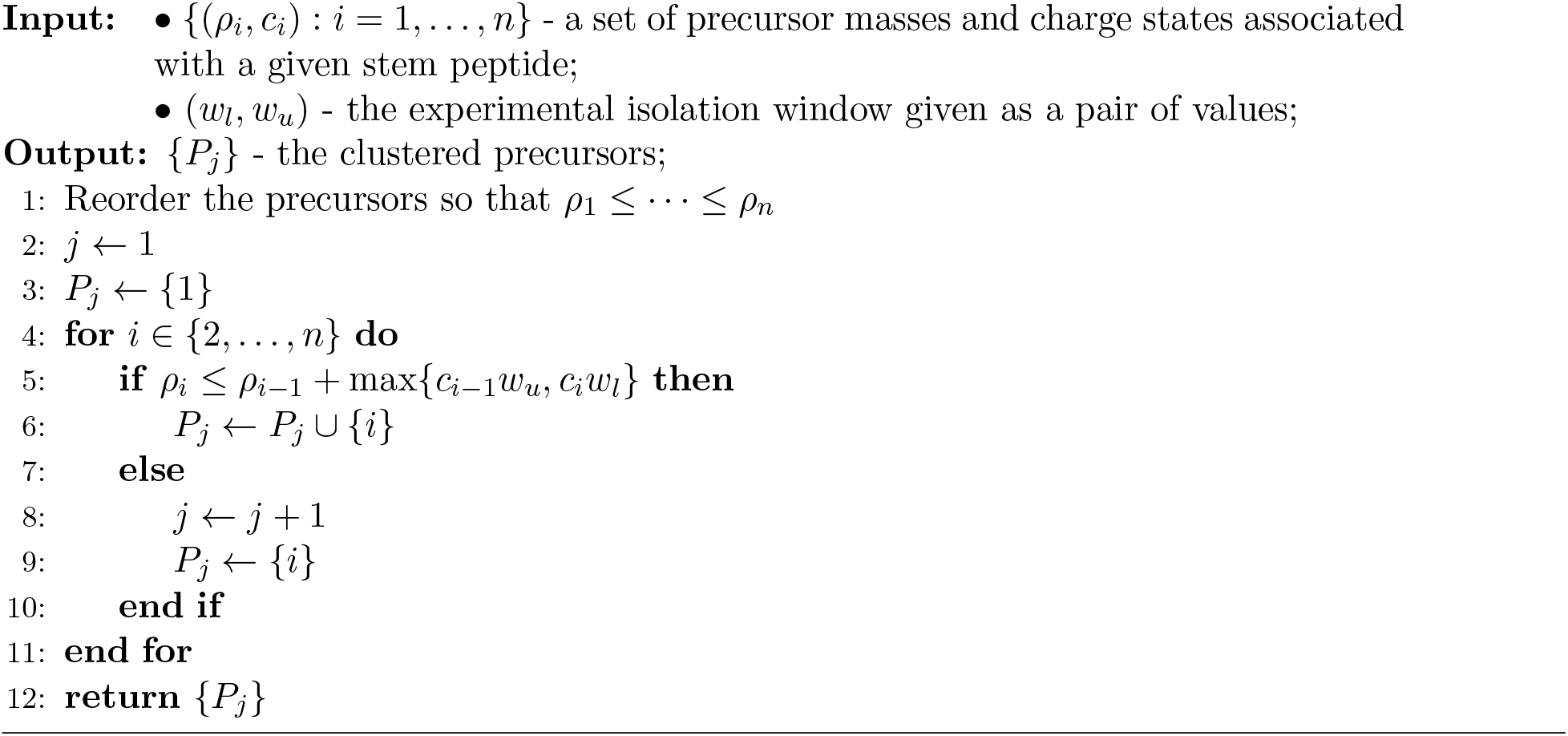

#### Algorithm 6

**Generating the auxiliary list of target PSMs with variants of the discovered peptides**

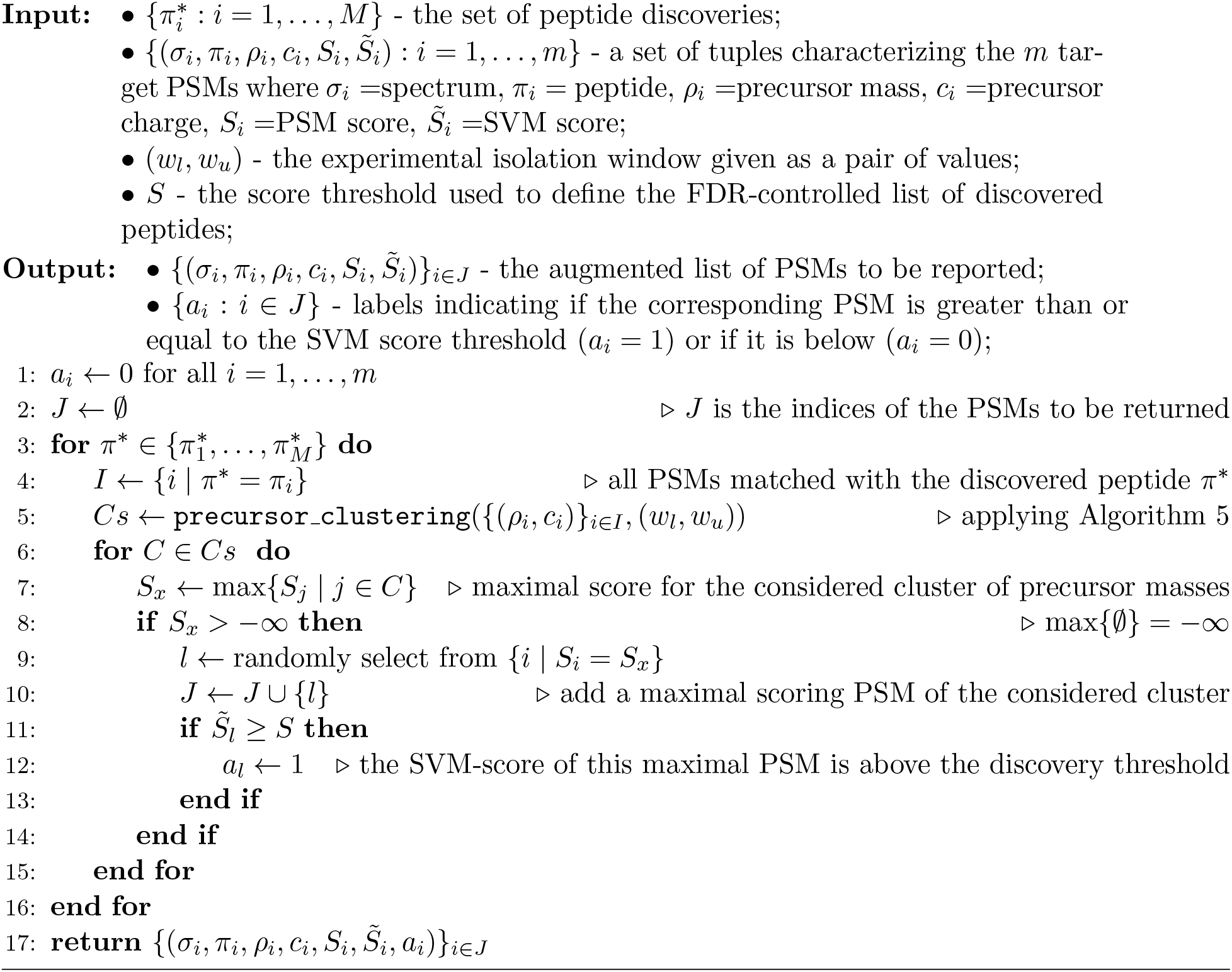

## References

[1] R. F. Barber and Emmanuel J. Candès. Controlling the false discovery rate via knockoffs. The Annals of Statistics, 43(5):2055–2085, 2015.

[2] Y. Danilova, A. Voronkova, P. Sulimov, and A. Kertész-Farkas. Bias in false discovery rate estimation in mass-spectrometry-based peptide identification. Journal of Proteome Research, 18(5):2354–2358, 2019.

[3] B. Diament and W. S. Noble. Faster SEQUEST searching for peptide identification from tandem mass spectra. Journal of Proteome Research, 10(9):3871–3879, 2011.

[4] J. E. Elias and S. P. Gygi. Target-decoy search strategy for increased confidence in large-scale protein identifications by mass spectrometry. Nature Methods, 4(3):207–214, 2007.

[5] J. Freestone, L. Käll, W. S. Noble, and U. Keich. Semi-supervised learning while controlling the fdr with an application to tandem mass spectrometry analysis. bioRxiv, 2023. https://www.biorxiv.org/content/10.1101/2023.10.26.564068v3.

[6] Jack Freestone, William S Noble, and Uri Keich. Analysis of tandem mass spectrometry data with conga: Combining open and narrow searches with group-wise analysis. Journal of Proteome Research, 23(6):1894–1906, 2024.

[7] Jack Freestone, William Stafford Noble, and Uri Keich. Reinvestigating the correctness of decoy-based false discovery rate control in proteomics tandem mass spectrometry. Journal of Proteome Research, 23(6):1907–1914, 2024.

[8] V. Granholm, W. S. Noble, and L. Käll. On using samples of known protein content to assess the statistical calibration of scores assigned to peptide-spectrum matches in shotgun proteomics. Journal of Proteome Research, 10(5):2671–2678, 2011.

[9] V. Granholm, W. S. Noble, and L. Käll. A cross-validation scheme for machine learning algorithms in shotgun proteomics. BMC Bioinformatics, 13(Suppl 16):S3, 2012.

[10] K. He, Y. Fu, W.-F. Zeng, L. Luo, H. Chi, C. Liu, L.-Y. Qing, R.-X. Sun, and S.-M. He. A theoretical foundation of the target-decoy search strategy for false discovery rate control in proteomics. arXiv, 2015. https://arxiv.org/abs/1501.00537.

[11] L. Käll, J. D. Canterbury, J. Weston, W. S. Noble, and Michael J MacCoss. Semisupervised learning for peptide identification from shotgun proteomics datasets. Nature Methods, 4(11):923–925, 2007.

[12] A. Keller, A. I. Nesvizhskii, E. Kolker, and R. Aebersold. Empirical statistical model to estimate the accuracy of peptide identification made by MS/MS and database search. Analytical Chemistry, 74:5383–5392, 2002.

[13] A. Kertesz-Farkas, F. L. Nii Adoquaye Acquaye, K. Bhimani, J. K. Eng, W. E. Fondrie, C. Grant, M. R. Hoopmann, A. Lin, Y. Y. Lu, R. L. Moritz, M. J. MacCoss, and W. S. Noble. The Crux Toolkit for Analysis of Bottom-Up Tandem Mass Spectrometry Proteomics Data. Journal of Proteome Research, 22(2):561–569, Feb 2023.

[14] J. Klimek, J. S. Eddes, L. Hohmann, J. Jackson, A. Peterson, S. Letarte, P. R. Gafken, J. E. Katz, P. Mallick, H. Lee, A. Schmidt, R. Ossola, J. K. Eng, R. Aebersold, and D. B. Martin. The standard protein mix database: a diverse data set to assist in the production of improved peptide and protein identification software tools. Journal of Proteome Research, 7(1):96–1003, 2008.

[15] A. T. Kong, F. V. Leprevost, D. M. Avtonomov, D. Mellacheruvu, and A. I. Nesvizhskii. MSFragger: ultrafast and comprehensive peptide identification in mass spectrometry-based proteomics. Nature Methods, 14(5):513–520, 2017.

[16] A. Lin, T. Short, W. S. Noble, and U. Keich. Improving peptide-level mass spectrometry analysis via double competition. Journal of Proteome Research, 21(10):2412–2420, 2022.

[17] L. Martens, H. Hermjakob, P. Jones, M. Adamsk, C. Taylor, D. States, K. Gevaert, J. Vandekerckhove, and R. Apweiler. PRIDE: The proteomics identifications database. Proteomics, 5(13):3537–3545, 2005.

[18] D. H. May, K. Tamura, and W. S. Noble. Detecting modifications in proteomics experiments with Param-Medic. Journal of Proteome Research, 18(4):1902–1906, 2019.

[19] P. Sulimov and A. Kertész-Farkas. Tailor: A nonparametric and rapid score calibration method for database search-based peptide identification in shotgun proteomics. Journal of Proteome Research, 19(4):1481–1490, 2020.

[20] Bo Wen, Jack Freestone, Michael Riffle, Michael J MacCoss, William S Noble, and Uri Keich. Assessment of false discovery rate control in tandem mass spectrometry analysis using entrapment. bioRxiv, pages 2024–06, 2024.

